# MicroHapDB: a portable and extensible database of all published microhaplotype marker and frequency data

**DOI:** 10.1101/2020.04.08.032052

**Authors:** Daniel Standage, Rebecca Mitchell

## Abstract

Microhaplotypes are the subject of significant interest in the forensics community as a promising multi-purpose forensic DNA marker for human identification. Microhaplotype markers are composed of multiple SNPs in close proximity, such that a single NGS read can simultaneously genotype the individual SNPs and phase them in aggregate to determine the associated donor haplotype. Abundant throughout the human genome, numerous recent studies have sought to discover and rank microhaplotype markers according to allelic diversity within and among populations. Microhaplotypes provide an appealing alternative to STR markers for human identification and mixture deconvolution, but can also be optimized for ancestry inference or combined with phenotype SNPs for prediction of externally visible characteristics in a multiplex NGS assay. Designing and evaluating panels of microhaplotypes is complicated by the lack of a convenient database of all published data, as well as the lack of population allele frequency data spanning disparate marker collections. We present MicroHapDB, a comprehensive database of published microhaplotype marker and frequency data, as a tool to advance the development of microhaplotype-based human forensics capabilities. We also present population allele frequencies derived from 26 global population samples for all microhaplotype markers published to date, facilitating the design and interpretation of custom multi-source panels. We submit MicroHapDB as a resource for community members engaged in marker discovery, population studies, assay development, and panel and kit design.

## 1 INTRODUCTION

Well-studied short tandem repeat (STR) markers have formed the basis of forensic human identification methods since the 1990s. The most common strategy in practice today utilizes several fluorescent dyes to type 20 or more STR markers in a single polymerase chain reaction (PCR) followed by capillary electrophoresis (CE) detection (Butler, 2010). The resulting DNA profiles, combined with STR allele frequency estimates, can then be used to calculate match statistics or evaluate the relative weight of evidence for competing propositions in a likelihood ratio framework (Butler, 2015; Cowell et al., 2015; Bleka et al., 2016a,b). Statistics obtained via STR typing can provide high confidence given the number of independent markers in an assay and the multiallelic nature of each marker.

Despite impressive recent improvements in DNA sequencing technologies, next-generation sequencing (NGS) assays of single nucleotide polymorphism (SNP) markers have seen slow adoption for forensic human identification. The ability to genotype sufficient numbers of SNPs to achieve suitable statistical power remains beyond the scope of many forensics laboratories. A relevant factor is the forensics community’s strong disinclination, on ethical and privacy grounds, to use DNA markers associated with human diseases or conditions, which limits the utilization of many commonly used microarrays and SNP chips. Also, because the majority of SNPs are bi-allelic, less population-level diversity is observed at each marker than at multi-allelic STRs, resulting in reduced discriminatory power when comparing reference and evidentiary samples. While this can be compensated for to some extent with a larger panel (SNPs are incredibly abundant in the human genome), the statistical requirement for markers that are inherited independently complicates panel design and places a practical limit on the resulting panel size.

*Microhaplotypes* (often abbreviated as *microhaps* or *MHs*) have recently prompted considerable interest in the forensics community as a promising alternative to independent SNPs and STRs for human identification (Kidd et al., 2018; Oldoni et al., 2019). A microhaplotype marker is defined by multiple SNPs^1^ residing within a short genomic distance whose state is reported as the allelic combination of all its component SNPs—that is, the haplotype. Here, “short” simply means a few hundred base pairs or fewer, ensuring a low frequency of recombination within the marker, and that a single NGS read or read pair can span all of the marker’s component SNPs. This length constraint enables each distinct read to both genotype and phase its target marker; that is, to determine 1) the individual allele of each component SNP, as well as 2) the haplotype. Even if a particular microhap is composed only of biallelic SNPs, the presence of multiple component SNPs makes it possible to observe several haplotypes at the marker, substantially increasing its discriminatory power over independent SNPs.

With a targeted NGS sequencing assay, a sufficient number of reads are collected to confidently genotype each marker, differentiating between true haplotypes and those arising from sequencing error. Microhap markers exhibit none of the stutter artifacts commonly observed in PCR-based STR assays, and the substitution and homopolymer errors common to some NGS platforms are easily resolved with sufficient depth of coverage. The restricted length of microhap markers makes them suitable for typing degraded samples, and the ability of NGS assays to capture additional rare SNPs within the microhaplotype can provide valuable information for improving mixture deconvolution compared to conventional CE-based STR assays.

Another notable benefit of microhaps is that they can be selected not only for high *within*-population variation, but also for high *between*-population variation, facilitating prediction of biogeographic ancestry (Oldoni et al., 2017; Zhu et al., 2019; Chen et al., 2019b). It is thus possible to design a comprehensive forensic panel using a combination of microhap and SNP markers that will enable identification, mixture detection and deconvolution (Bennett et al., 2019), ancestry inference, and prediction of externally visible characteristics (Walsh et al., 2013; Ruiz et al., 2013; Crawford et al., 2017) in a single NGS-based assay.

Microhaps are abundant in the human genome, and thus discovering and ranking them for different purposes is an area of active research interest in the forensics community. In just the last few years, numerous studies presenting new microhap marker collections have been published, together totaling more than 400 markers (Hiroaki et al., 2015; Kidd et al., 2018; van der Gaag et al., 2018; Voskoboinik et al., 2018; Staadig and Tillmar, 2019; Chen et al., 2019a; de la Puente et al., 2020). While two of these studies also include allele frequency data for multiple population samples—including 83 populations^2^ in Kidd et al. (2018) and 3 populations in van der Gaag et al. (2018)—the others present either no frequency data or data for a single population sample, with little overlap between studies. The absence of a single point of access for published microhaps and the paucity of allele frequency data spanning disparate data sets are obstacles to developing and evaluating custom panels composed of microhap markers selected from different published collections.

To support the development of microhaplotype-based human forensics capabilities, we have compiled a database of all published microhap marker definitions and allele frequencies. MicroHapDB is a portable database that, once installed, can be accessed by the user without an Internet connection. The entire contents of the database are distributed with each copy of MicroHapDB, and instructions for adding private data to a local instance of the database are provided. The same instructions can alternatively be used by MicroHapDB maintainers or interested community contributors to submit new markers and allele frequencies for review and potential inclusion in the public database. MicroHapDB is designed to be user-friendly both for forensic practitioners and researchers, and supports a variety of access methods including browsing, simple or complex text queries, and programmatic database access via a Python application programming interface (API). Finally, to increase the value of the published microhaps in aggregate, we have used 2,504 fully phased genomes from the 1000 Genomes Project (Auton et al., 2015) to estimate allele frequencies in 26 global populations for 412 microhap markers. MicroHapDB is a valuable resource for researchers, practitioners, and commercial entities engaged in marker discovery, population studies, assay development and validation, and design of custom panels and kits for forensic applications.

## 2 MATERIALS AND METHODS

### 2.1 Database Design

The contents of the MicroHapDB database are stored in seven tables, distributed as plain-text tab-delimited files. Three of the tables constitute the “core database,” and include population sample descriptions, marker definitions, and microhaplotype frequencies (Figure 1). The remaining four tables contain ancillary metadata: a cross reference of third-party identifiers to MicroHapDB identifiers; data for a small number of indel variants present in the database; the genomic sequences spanning each marker (to facilitate amplicon design); and a mapping of variant identifiers (rsIDs) to marker names.

**Figure 1.**
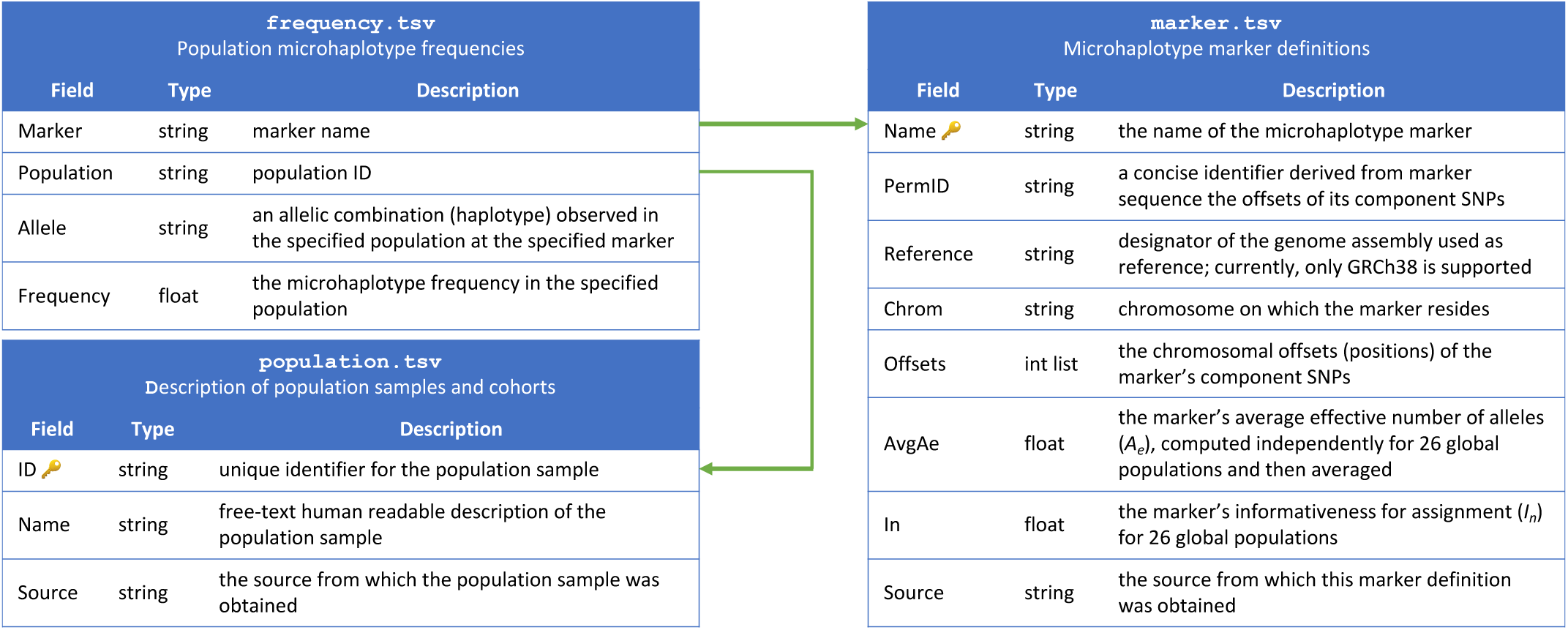
Schema for the MicroHapDB core database.

### 2.2 User Interface

The primary interface for querying MicroHapDB is the UNIX command line. The `microhapdb` command provides three operations corresponding to the three tables in the core database: `microhapdb marker`, `microhapdb population`, and `microhapdb frequency`. Executing any of these commands with no additional arguments will print the entire contents of the specified table to the terminal for browsing. Each command also enables a user to restrict the printed results to data matching a particular identifier, source, or genomic region.

By default, all results are printed in tabular format. Population data can optionally be printed in a “detail” format summarizing the number of markers for which microhaplotype frequency is available and the total number of microhaplotypes (microhaps) observed in the population sample. Marker data can optionally be printed in FASTA format for use with third-party programs, or in a “detail” format showing the genomic location of the marker and its component SNPs, the core marker sequence (spanning only the microhap’s most distal component SNPs), all haplotypes observed at the marker, and a candidate amplicon sequence for the marker (for which the amount of flanking sequence can be configured using the `--delta` and `--min-length` parameters): see Figure 2.

**Figure 2.**
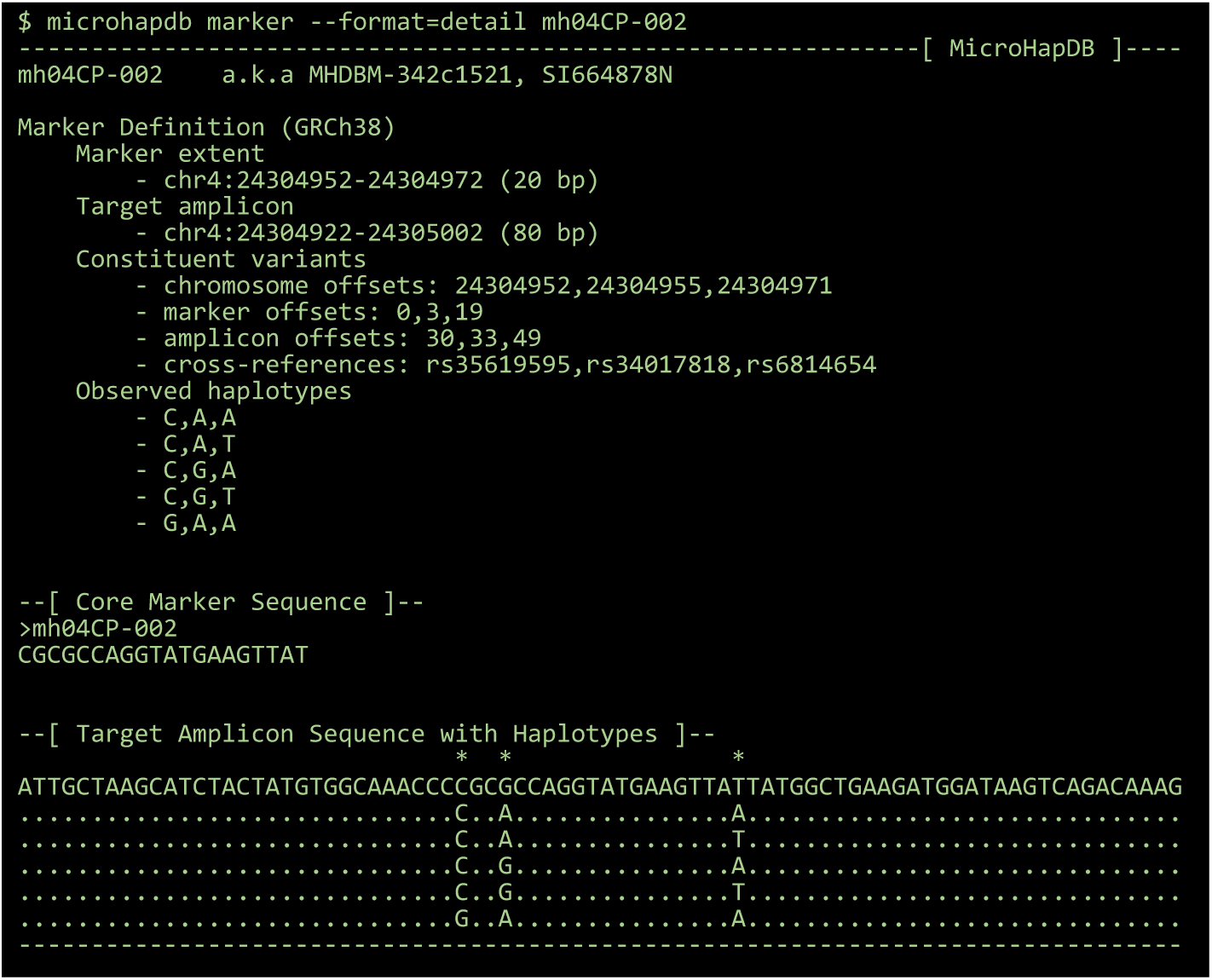
The “detail” view for a 3-SNP microhaplotype marker, displayed using the MicroHapDB command-line interface.

Once MicroHapDB is installed on a computer, all database list and search operations access only the data files resident on that machine. MicroHapDB doesn’t require or permit requests to transfer data to or from databases residing on remote machines.

Installed as a Python package, MicroHapDB also supports programmatic access to the core and ancillary database tables. After invoking `import microhapdb`, users can write custom code to query and analyze marker, population, or frequency data resident in the database tables, pre-loaded into memory as pandas dataframes (McKinney, 2011). Alternatively, users can execute the `microhapdb --files` command from the UNIX shell to show the location of the database table files, which can be imported directly into R, Excel, or any other data analytics environment preferred by the user.

### 2.3 Implementation and Availability

At its core, MicroHapDB is composed of a small number of tabular plain-text data files containing marker, population, and frequency information for published microhaps. These files enclose the entire contents of the database. In contrast to many databases of genetic variation, each instance of MicroHapDB stores the entire contents of the database locally. MicroHapDB does not communicate with any central database server, and network connections are only used to install the database or upgrade to a newer version.

The user interface described in Section 2.2 is implemented in a Python package that can be installed and upgraded using the Bioconda software manager (Grϋning et al., 2018). Source code for MicroHapDB is published open-access on the GitHub platform at https://github.com/bioforensics/MicroHapDB/ and is free for commercial and non-commercial use under a permissive open source license. The authors operate the MicroHapDB project under an open governance model that facilitates and encourages contributions from the community.

The software and procedure used by the authors to build the database is also published on the MicroHapDB GitHub repository. During the build process, data from several sources is independently pre-processed and standardized, and then all sources are aggregated and sorted to compile the final database. This strategy, described further in Section 2.4, serves several purposes. First, it provides a clear mechanism for the authors or other community members to extend the database in the future as additional marker and frequency data is published in the literature. Second, the same mechanism enables interested users to supplement the public MicroHapDB database with private data in a safe and secure way. By following the guidelines in the database build instructions provided in the MicroHapDB repository, a user can rebuild their local copy of MicroHapDB with additional sources of marker and/or frequency data. Because MicroHapDB doesn’t communicate with any central database, changes made to a user’s local copy of MicroHapDB do not propagate to GitHub or any other location. Third, it permits careful scrutiny of the entire database construction process by any interested party in case errors in the database contents are ever discovered.

### 2.4 Data Collection and Pre-processing

MicroHapDB was compiled from seven distinct sources, each of which organized and reported data in a unique format. Extracting the relevant data and cross-referencing with public databases of genomic variation required a combination of manual and automated strategies uniquely designed for each distinct source. The result of this preliminary data acquisition and pre-processing was a collection of seven data sets with consistently formatted population descriptions, marker definitions, and allele frequencies. Once data from each distinct source was collected, cross-referenced with the GRCh38 human reference genome, and standardized, the final database compilation was performed by aggregating and sorting all data sources.

Source code and corresponding technical documentation describing data collection, pre-processing, and aggregation of the final database is available at https://github.com/bioforensics/MicroHapDB/tree/0.5/dbbuild.

### 2.5 Estimation of Haplotype Frequencies for 26 Global Populations

MicroHapDB includes data from several distinct sources, but the availability of population frequencies for published microhaps is inconsistent. For some markers, frequencies are reported for dozens of population samples. Other markers have frequencies reported only for a single cohort, and yet other markers have no frequencies reported whatsoever. Designing and testing panels composed of markers from multiple distinct sources is possible, but prior to the release of MicroHapDB, interpretation of any sample assayed using such a panel would require the development of appropriate frequency data. We used 2,504 fully phased genotypes from a publicly available large-cohort study (Auton et al., 2015) to estimate population frequencies for all published microhaps across a set of 26 global population samples. These frequencies were recently published in MicroHapDB version 0.5.

MicroHapDB version 0.5 contains definitions for 417 microhap markers. Five of these markers^3^ are defined by rare variants not genotyped in the 1000 Genomes Project Phase 3 data, and were thus excluded from this analysis. For each of the remaining 412 markers, population frequencies for 26 global populations were estimated using the following procedure. First, phased genotype records for each of the marker’s component variants were retrieved using the variants’ rsIDs. Next, the phased genotypes were aggregated to determine the two haplotypes for each individual at the marker (or the single haplotype observed at X chromosome markers in males). Then, noting the population sample with which each individual was associated, a tally of haplotypes was compiled for each population. Finally, the haplotype tallies for each population sample were normalized by the corresponding number of alleles to compute the final frequency estimates.

### 2.6 Calculation of Measures of Variation Within and Among Populations

Microhaps are suitable for numerous forensic applications. Two common statistics used for ranking microhaps are the effective number of alleles *A*_*e*_ and the informativeness for assignment *I*_*n*_ (Crow and Kimura, 1970; Rosenberg et al., 2003; Kidd and Speed, 2015; Kidd et al., 2018). The *A*_*e*_ statistic is the reciprocal of a marker’s homozygosity. For a marker with *N* alleles, *A*_*e*_ is computed as

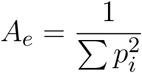

where *p*_*i*_ is the frequency of allele *i* ∊ *N* and summation is over all alleles. It is a measure of allelic variation within a population, and corresponds to a marker’s power for individual identification. For a microhaplotype with *N* SNPs, the maximal *A*_*e*_ value of 4^*N*^ occurs if and only if every possible allelic combination is observed at equal frequencies in the population. In reality, only a subset of possible allelic combinations are generally present in a population, and typically at unequal frequencies, resulting in *A*_*e*_ values that most commonly fall between 1.5 and 4.5 for previously reported microhap markers. The minimal *A*_*e*_ value of 1 occurs when only a single haplotype is observed at the locus in a population.

By contrast, *I* _*n*_ measures the extent of population-specific allelic variation among a set of populations, and corresponds to a microhap’s power for predicting an individual’s biogeographic ancestry. The *I*_*n*_ statistic for a marker with *N* alleles across *K* populations is calculated as

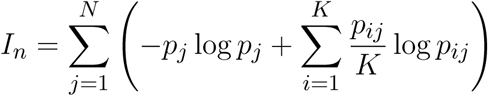

where *p*_*ij*_ is the frequency of allele *j* in population *i*, log is the natural log, and 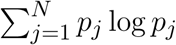 is the expected log-likelihood associated with drawing an allele from a hypothetical “average” population (frequencies averaged over all *K* populations). The minimal *I*_*n*_ of 0 occurs when all alleles have equal frequencies in all populations, and the maximal value log *K* occurs when *N* ≥ *K* and no allele is found in more than one population (Rosenberg et al., 2003).

As a final post-processing step in the MicroHapDB database build procedure, *A*_*e*_ and *I*_*n*_ statistics were computed for all markers in MicroHapDB.^4^ Using population microhaplotype frequencies computed from the 1000 Genomes Project Phase 3 genotypes (Auton et al., 2015), MicroHapDB scripts computed per-marker *A*_*e*_ values individually for each population, and then the arithmetic mean over all 26 populations. The result is listed in the `AvgAe` column of MicroHapDB’s `markers` table. *I*_*n*_ statistics over 26 populations were calculated with the same frequency data using INFOCALC (https://rosenberglab.stanford.edu/infocalc.html), and are listed in the `In` column of the `markers` table.

## 3 RESULTS

### 3.1 MicroHapDB Aggregates Data for More than 400 Microhaplotypes, Including Measures of Within- and Between-population Variation

MicroHapDB version 0.5, released in February 2020, includes descriptions of 102 cohorts and population samples from six sources, 417 marker definitions from seven sources, and numerous population frequencies for 5,373 observed haplotypes from six sources, all together comprising a total of 113,995 records. This database represents a comprehensive collection of all microhaplotype (microhap) data published to date.

The number of single nucleotide polymorphisms (SNPs) used to define microhaps ranges from 2 to 49, with an average of 3.77 SNPs per marker. MicroHapDB includes 115 markers defined by two SNPs, 171 defined by three SNPs, 87 defined by four SNPs, 20 defined by five SNPs, 5 defined by six SNPs, and 19 defined by seven or more SNPs.

Microhap markers are defined on all autosomes as well as the X chromosome. Forty-five marker definitions overlap with other markers. Most of these (40/45) were defined by Staadig and Tillmar (2019), which includes 11 exact duplicates and 29 probable adjustments to markers from other sources. The distance between each marker and the closest non-overlapping marker ranges between 143 bp and more than 56 Mb. Out of 417 markers in MicroHapDB, 152 (36.5%) reside within 1 Mb of their closest neighbor, and 53 (12.7%) reside within 100 kb of their closest neighbor (Figure 3A). Any set of markers separated by a small physical distance is likely in high linkage disequilibrium (LD) and the individual markers would therefore not be independent. DNA profiles containing information for linked markers further complicates interpretation. This requires either sophisticated statistical modeling to account for the dependencies between markers or adopting an either/or strategy in which one of the loci is discarded when both produce reliable data.

**Figure 3.**
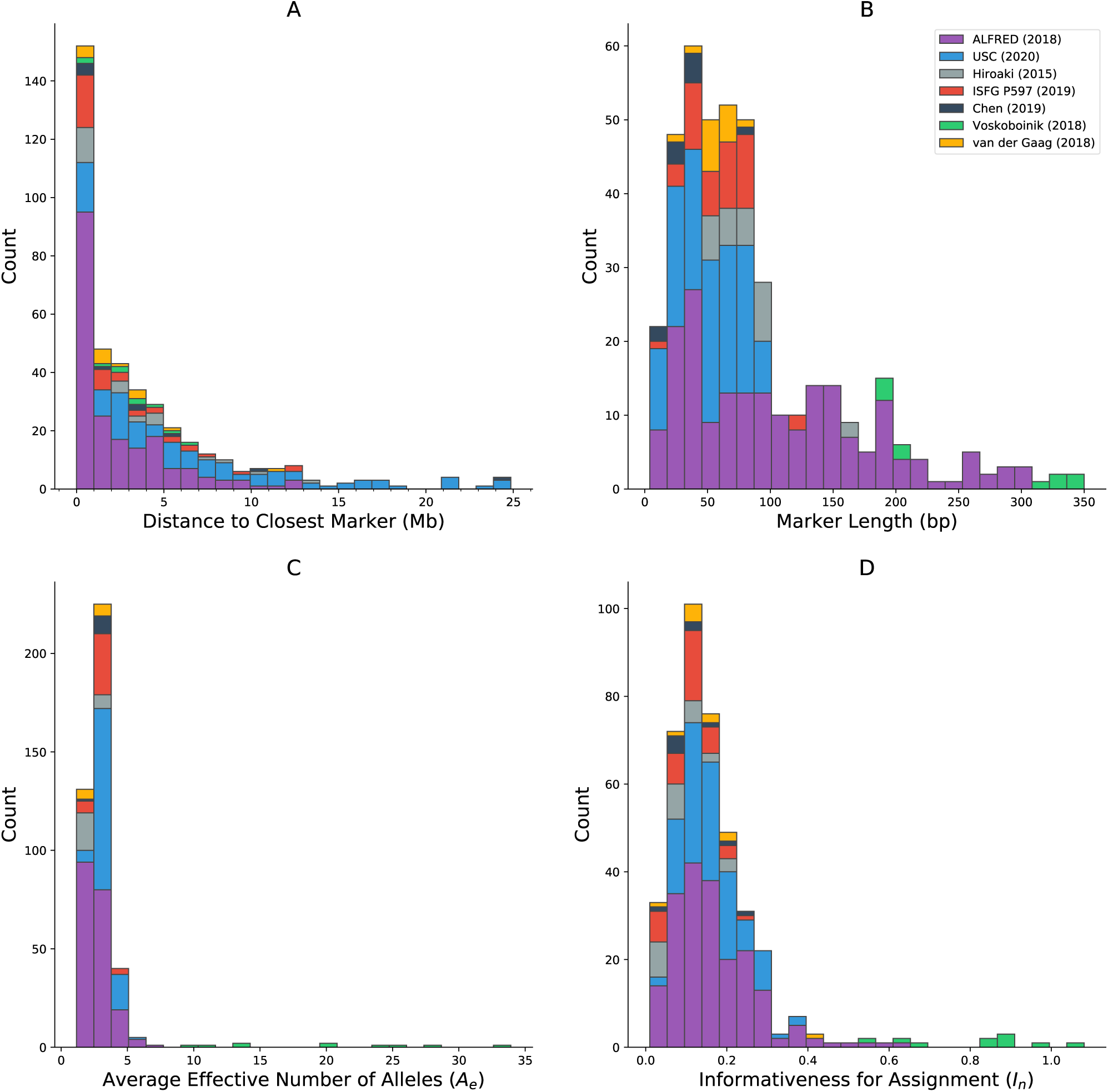
Histograms showing the distribution of various characteristics of microhaplotype markers in MicroHapDB, shaded by publication source: (A) distance between a marker and its closest non-overlapping marker (outliers not shown: 42.8 Mb and 56.1 Mb); (B) marker length; (C) effective number of alleles [*A*_*e*_], averaged over 26 global populations; (D) informativeness for assignment [*I*_*n*_] to 26 global populations.

Published microhaps occupy a wide range of lengths, with core marker length (the number of nucleotides spanning the most distal SNPs that define the marker, inclusive) ranging from 4 bp to 350 bp (Figure 3B). The majority of the microhaps in MicroHapDB (307/417; 73.6%) span less than 100 bp, and a substantial minority (145/417; 34.8%) span less than 50 bp.

The per-marker effective number of alleles (*A*_*e*_) was computed independently for 26 global populations, and then averaged across all populations (Materials and Methods). This statistic serves as a measure of within-population allelic diversity observed at a particular marker, and corresponds to the marker’s power for individual identification. A previous study (Kidd and Speed, 2015) proposed an *A*_*e*_ threshold of 3, above which a microhaplotype can be considered “exceedingly useful” for forensic purposes. Average *A*_*e*_ values for microhaps in MicroHapDB range from 1.16 to 33.92, with a mean of 3.28 (Figure 3C). Most markers in MicroHapDB (238 / 412, 57.8%) have an average *A*_*e*_ below 3, and only 17 markers (4.1%) have an average *A*_*e*_ above 5.

Marker informativeness for assignment (*I*_*n*_) was computed for the same 26 global populations (Materials and Methods). This statistic serves as a measure of variation among populations, and corresponds to the marker’s power for predicting an individual’s biogeographic ancestry. *I*_*n*_ values for markers in MicroHapDB fall between 0.01 and 1.08, with a mean of 0.17 (Figure 3D). Eight markers have an *I*_*n*_ value greater than 0.682, the highest *I*_*n*_ value previously reported for a microhap (Kidd et al., 2018)—we note however that these *I*_*n*_ values were computed for a different set of populations and are therefore not directly comparable.

Ten highly polymorphic microhaps reported by Voskoboinik et al. (2018) stand out from microhaps published in other sources in several ways. Originally defined by locus boundaries rather than by explicit lists of SNP identifiers, these 10 microhaps include more SNPs (14-49 per marker) than any other marker in MicroHapDB. They have the highest average *A*_*e*_ values in MicroHapDB by a significant margin, and 9 of these markers are included in MicroHapDB’s top 10 microhaps ranked by *I*_*n*_. It is worth noting that these microhaps are also among the longest, reflecting the study’s distinct selection criteria and sequencing and evaluation strategy. The longest 5 markers in this set are also the 5 longest markers in MicroHapDB, and the remaining 5 are above the 89th percentile in length with respect to all markers in MicroHapDB.

### 3.2 Microhaplotype Frequencies for 26 Global Populations Enable Interpretation of Multi-source Panels

Interpretation of any microhap typing result requires the use of appropriate microhaplotype frequency data. Prior to the release of MicroHapDB, availability of frequency data was inconsistent for published microhaps, with some sources providing frequencies for numerous population samples, while other sources providing frequencies for only a single population sample, or no frequency data at all.

MicroHapDB provides a comprehensive set of frequencies for all microhap markers to date. Estimated using 2,504 fully phased genotypes from Phase 3 of the 1000 Genomes Project (Auton et al., 2015), the MicroHapDB database contains frequencies for 412 microhaplotype markers across 26 global population samples. A total of 113,477 frequencies for 5,373 alleles furnish a broad view of marker variation amongst Africans, admixed Americans, East Asians, Europeans, and South Asians.

## 4 DISCUSSION

In this study, we report the development of a comprehensive database of published microhaplotype (microhap) marker and frequency data. We describe the estimation of microhap frequencies in 26 global population samples, and the use of these frequencies to compute measures of allelic variation, enabling the ranking of microhaps for different forensic applications. This extensive collection of allele frequencies and ranking statistics will facilitate the design and interpretation of forensic panels that include markers from distinct sources without the need for extensive development of frequency data up front. MicroHapDB is a free open-access resource that contains information for all microhaps published as of February 2020. It requires minimal computing resources to install and maintain, and is designed to be easily extended with additional sources of public or private microhap data in the future. We hope MicroHapDB will democratize and accelerate advances in microhap-based forensics capabilities by enabling researchers and companies to focus solely on marker discovery, or solely on population surveys, or solely on panel and kit design, without the need to invest in development of all of the above.

MicroHapDB provides *A*_*e*_ and *I*_*n*_ values for ranking microhaps for different forensic purposes. The genomic coordinates of each marker are also stored in MicroHapDB, enabling convenient calculation of physical distances between markers residing on the same chromosome. However, in addition to normal considerations that must always be addressed when designing a forensic DNA panel (e.g. amplicon sizes, primer kinetics, off-target amplification), correct interpretation of DNA profiles requires researchers to determine the independence of markers in a proposed panel based on the extent of linkage between the markers in the population(s) of interest. The length of candidate markers is also an important consideration depending on the sequencing technology utilized and the priority of recovering profiles from low input or low quality DNA samples.

Unlike conventional short tandem repeat (STR) markers, no comprehensive database analogous to the FBI’s Combined DNA Index System (CODIS) databases (https://www.fbi.gov/services/laboratory/biometric-analysis/codis) yet exists for microhap profiles. Constructing, populating, and performing requisite quality control for such a database will require substantial time and investment, which likely could only be pursued in earnest as the forensic community approaches consensus regarding optimal targets and assays. We expect that MicroHapDB can play a useful role in that process.

A major challenge in establishing a database like MicroHapDB, or a shared national database of microhap profiles, or indeed in communicating clearly about microhaps in the scientific literature, is the lack of consistency in nomenclature and in the way that markers are defined. A few papers describing microhap markers have used the nomenclature proposed by Kidd (2016), with marker names such as `mh01KK-172`, `mh01CP-007`, and `mh06PK-25713`. Other papers used a variety of *ad hoc* marker designators, such as `1` and `MH02`. MicroHapDB has adopted the Kidd nomenclature since its inception in 2018, and for sake of consistency has applied it to microhap collections where it was not previously used (e.g. `mh01NH-01` for `1` and `mh01AT-02` for `MH02`).

The question of how microhap markers are *defined* is at least as consequential as how they are *labeled*. A small number of published microhaps have been defined as a specific (but undisclosed) set of single nucleotide polymorphisms (SNPs) at a genomic locus, the endpoints of which are indicated using coordinates on a reference genome assembly. Other microhaps are defined by a designator that refers to a set of SNPs at a particular locus, but whose specific component SNPs have been adjusted over time to improve the marker’s performance. Ambiguous marker definitions of these kinds create substantial challenges for reproducibility and establishing provenance, and should be avoided. de la Puente, Phillips, et al. (2020) propose that microhaps should forgo marker names altogether (e.g. `mh01USC-1pA` or `1pA`) in favor of an explicit list of SNP variants, as designated by dbSNP rsIDs (e.g. `rs28503881,rs4648788,rs72634811,rs28689700`). We strongly endorse this sentiment, although we concede the convenience of concise marker designators, especially when marker definitions are composed of dozens of SNP variants. What is most critical is the need for marker definitions to be unambiguous and invariant over time.

This discussion highlights the tension that has been provoked by the emergence of NGS technologies in forensics. Conventional assays have required the design of probes for specific SNP targets, which are often genotyped independently and then phased statistically. In contrast, NGS assays permit recovery of the entire sequence at a microhap locus and the simultaneous genotyping and phasing of all its component SNPs, and indeed any additional intermediate (and often rare) SNPs. We anticipate that as NGS forensic assays become more routine, typing results for microhap assays will include all of the variants occurring in the sequenced genomic segment. This kind of typing result would have full “backwards compatibility” in that it could be used to determine the haplotype of *any* microhap marker explicitly defined at the locus. At the same time, full-coverage sequences of microhap loci will enable significant improvements in, e.g., mixture detection and deconvolution.

The question remains as to whether microhap designators should refer to *markers* (i.e., explicit sets of variants) or to *loci*. We suggest that minor addenda to the nomenclature proposed by Kidd (2016), such as the use of version numbers or other suffixes, would enable support for both. In the mean time, MicroHapDB searches based on genomic coordinates provide a convenient way to resolve spatial relationships between distinct marker definitions.

## AUTHOR CONTRIBUTIONS

DS conceived the study, constructed the database, implemented the software interface, and wrote the manuscript. DS and RM collected data, performed quality control, and edited and approved the manuscript.

## ACKNOWLEDGMENTS

We express gratitude to Kenneth Kidd, Christopher Phillips, and Lev Voskoboinik for responding to inquiries about published data sets, and for insightful discussions about microhaplotypes. We also thank Rebecca Just, Tim Stockwell, M.J. Rosovitz, and Adam Bazinet for their valuable feedback on drafts of this manuscript and early versions of the database.

## DATA AVAILABILITY STATEMENT

MicroHapDB version 0.5 has been archived in the Open Science Framework repository and is available at https://osf.io/fduza/. The MicroHapDB database and software can be installed and updated from the Bioconda repository: see https://bioconda.github.io/ for more details.

## Footnotes

1 While the vast majority of published microhaplotype markers are composed exclusively of SNPs, a small number include one or more insertion/deletion polymorphisms (indels).

2 As of the latest December 2018 update, there are now 96 populations in the Allele Frequency Database (ALFRED) for which microhaplotype frequency data is available.

3 mh06PK-24844, mh0XUSC-XqD, mh11PK-63643, mh15PK-75170, and mh22PK-104638

4 With the exception of the 5 markers discussed in footnote 3.

